# Resource misallocation as a mediator of fitness costs in antibiotic resistance

**DOI:** 10.1101/456434

**Authors:** Andrej Trauner, Amir Banaei-Esfahani, Sebastian M. Gygli, Philipp Warmer, Julia Feldmann, Seyedehsara Shafieechashmi, Katja Eschbach, Mattia Zampieri, Sonia Borrell, Ben C. Collins, Christian Beisel, Ruedi Aebersold, Sebastien Gagneux

## Abstract

Antimicrobial resistance poses a threat to global health and the economy. It is widely accepted that, in the absence of antibiotics, drug resistance mutations carry a fitness cost. In the case of rifampicin resistance in fast-growing bacteria, this cost stems from a reduced transcription rate of the RNA polymerase resulting in slower ribosome biosynthesis. However, this relationship does not apply in the slow-growing *Mycobacterium tuberculosis*, where the true mechanism of fitness cost of rifampicin resistance as well as the impact of compensatory evolution remain unknown. Here we show, using global transcriptomic and proteomic profiling of selected *M. tuberculosis* mutants and clinical strains, that the fitness cost of rifampicin resistance in *M. tuberculosis* is the result of the physiological burden caused by aberrant gene expression. We further show that the perceived burden can be increased, effectively suppressing the emergence of drug resistance.

---

Antimicrobials are one of the cornerstones of modern medicine (Laxminarayan et al., 2016). The global increase of antimicrobial resistance (AMR) poses an existential threat, claiming an increasing number of lives and resources (O’Neill, 2016). We currently have access to a wide array of antibiotics, but their efficacy is waning, making safeguarding existing and future drugs a high priority. Understanding the mechanisms and drivers of AMR (Holmes et al., 2016), including the underlying biology, will be key to that process.

Antibiotics target essential bacterial processes. Modification of their targets is an important mechanism through which AMR emerges. It is therefore not surprising that AMR often comes with a fitness cost (Melnyk et al., 2015). Fitness cost is a broad concept capturing any negative deviation in the proliferation of a mutant from its ancestor: for example, a decreased growth rate *in vitro,* or in the case of pathogens, a decreased ability to transmit or cause disease. The physiological basis for the cost of drug resistance seems to be dependent on the antibiotic, bacterial species and environment (Andersson and Hughes, 2008) and is thus often unknown and likely to be multifaceted. One of the better studied examples is the cost of rifampicin resistance. Rifampicin targets the bacterial RNA polymerase (RNAP), and resistance to rifampicin is usually mediated by mutations in the β subunit of RNAP (Campbell et al., 2001). Several studies point to the rate of transcription, particularly as it pertains to the synthesis of ribosomal RNA and ribosomal proteins, as an important mediator of growth rate (Gourse et al., 1996; Thiele et al., 2009). A slowing down of transcription is therefore the prime mechanistic candidate for the cost of rifampicin resistance (Qi et al., 2014; Reynolds, 2000). The mechanism linking RNAP activity to ribosome biosynthesis provides a compelling explanation for the cost of rifampicin resistance in rapidly dividing bacteria such as *Escherichia coli* and *Pseudomonas aeruginosa* whose growth relies on the rapid replenishment of biosynthetic machinery lost through cell division (Ehrenberg et al., 2013). Importantly, the fitness cost of rifampicin resistance can be mitigated or even reversed through the acquisition of secondary, compensatory mutations in the α, β and β’ subunits of RNAP that seem to restore normal enzyme function (Qi et al., 2014; Song et al., 2014; Stefan et al., 2018).

Rifampicin-resistant *Mtb* is one of the major causes of AMR-associated mortality globally, claiming an estimated 240,000 lives in 2016 (WHO, 2017), and unlike in fast-growing bacteria, the rate of transcription does not seem to reflect the fitness cost of key *rpoB* mutations, measured either as growth rate *in vitro* or prevalence in the clinic (Gagneux et al., 2006; Gygli et al., 2017; Stefan et al., 2018). While relative fitness does seem to determine the clinical success of rifampicin-resistant *Mtb* (Grandjean et al., 2015), and compensatory mutations are frequently found in settings with a high burden of drug resistant TB (Casali et al., 2014; Comas et al., 2012; de Vos et al., 2013; Farhat et al., 2013), the basis for the fitness cost of rifampicin resistance remains unknown in *Mtb*. Understanding the mechanism by which *rpoB* mutations impair normal *Mtb* physiology could help identify new intervention points, through which we could stem the tide of existing and emergent rifampicin resistance.

We used the known ability of mutations in the beta barrel double Ψ (BBDP) domain of the β’ subunit of RNAP to compensate for the fitness cost of resistance mutations occurring in the β subunit in *Mtb* as a starting point (Molodtsov et al., 2017; Song et al., 2014; Stefan et al., 2018). Compensatory mutations improve patient to patient transmission of rifampicin-resistant strains (de Vos et al., 2013), and partially reverse biochemical changes imparted on RNAP by rifampicin-resistance mutations (Song et al., 2014; Stefan et al., 2018). We hypothesise that the same would be true for gene expression differences. Leveraging the knowledge of the role of RpoC mutations, we used transcriptomic and proteomic expression profiling to identify the signature of compensation and therefore infer the likely mediators of fitness cost in a collection of strains derived from a drug-susceptible clinical isolate (see Figure 1). Our findings point to the idiosyncratic consequences of expressional dysregulation as a key factor conferring a fitness cost to rifampicin resistance in *Mtb*. We expanded on this observation by profiling the expression signature of rifampicin resistance in a panel of genetically diverse clinical isolates sharing the same rifampicin resistance-conferring mutation: RpoB Ser450Leu. While we found very little evidence for a shared expression signature of rifampicin-resistance across the tested strain pairs, we show a correlation between the fitness cost of the rifampicin-resistance conferring mutation and the extent to which its presence imparts a deviation from the proteome composition of the wild-type. Finally, we show that this correlation could be exploited to suppress the emergence of rifampicin resistance.

**Figure 1:**
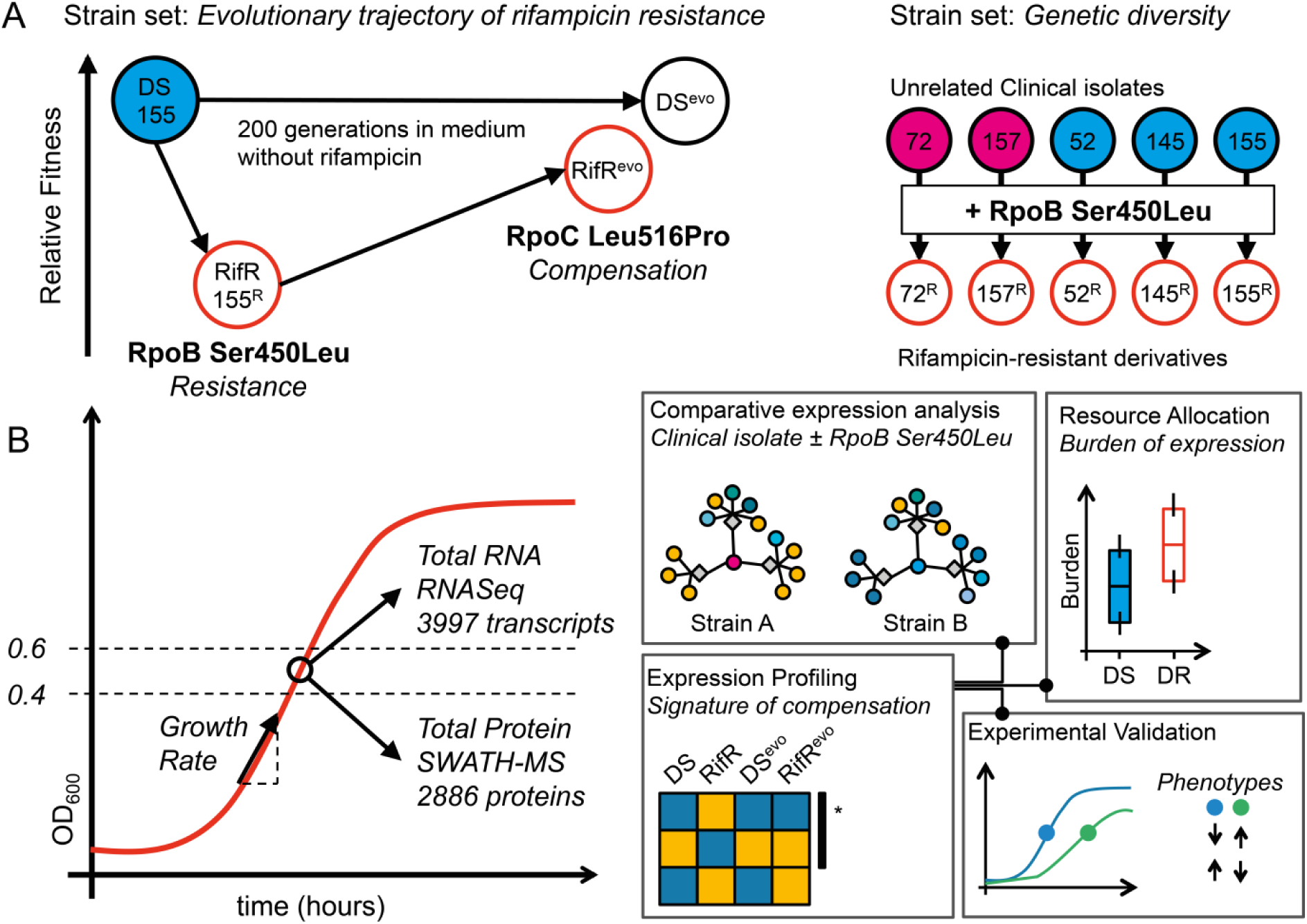
Conceptual workflow. **A.** Two complementary strain sets used for the experiments. Strains comprised in the “Evolutionary trajectory of rifampicin resistance” set were derived from a single clinical isolate (DS, N0155) by isolation of a Ser450Leu mutant in the lab and the subsequent passage for 200 generations in the absence of rifampicin. These strains were used to identify expression changes that are reversed by compensation -**signature of compensation**. The generalizability of our finding was checked using the “Genetic diversity strain set” containing five independent clinical isolates and their rifampicin-resistant derivatives. All rifampicin resistant strains shared the same resistance mutation – RpoB Ser450Leu. **B.** Experimental outline for the sampling and analyses.

## Results

### Compensatory mutations mitigate resistance-imposed expression changes

Physiological changes incurred by a fitness cost are likely to manifest as deviations in gene expression. Since mutations in the BBDP domain of the β’ subunit of RNAP mitigate the fitness cost of rifampicin-resistance mutations in *Mtb* (Molodtsov et al., 2017; Song et al., 2014; Stefan et al., 2018) they should also impact and therefore highlight expression changes that are relevant to the understanding of fitness cost of rifampicin resistance.

We previously reported the result of a directed evolution experiment in which we identified a mutation in the BBDP domain: RpoC Leu516Pro as a putative compensatory mechanism for the fitness cost of the rifampicin-resistance conferring mutation RpoB Ser450Leu in a clinical isolate(Comas et al., 2012). The strains generated by that study comprise the original drug-susceptible isolate (DS), its laboratory-derived rifampicin-resistant mutant (RpoB Ser450Leu, RifR) and the resulting evolved strains obtained by serial passage in the absence of rifampicin for 200 generations (DS^evo^and RifR^evo^, respectively, see Figure 1A). Together these strains offer a representative snapshot of the evolutionary process that passes through the initial emergence of (costly) drug resistance and leads to the establishment of a mature drug-resistant strain whose fitness is indistinguishable from its drug susceptible ancestor. We therefore hypothesised that comparative transcriptomic and proteomic expression profiling of these strains will allow us to determine the signature of the fitness cost associated with rifampicin resistance.

First, we determined the relative fitness of RifR. Using a mixed effect linear regression model to analyse growth assays, we noted a 26.4% decrease (CI^95%^: 21.5 – 31.0%, p < 0.001) in the growth rate of RifR when compared to DS. The comparison of their evolved counterparts – DS^evo^and RifR^evo^– showed no significant differences (−1.2%, CI^95%^: -10.8 – 7.1%, p = 0.814), illustrating the fact that RpoC Leu516Pro does indeed compensate the fitness cost of rifampicin resistance.

We aimed to identify differences in the baseline, unperturbed, gene expression as a proxy for describing the biological basis for reduced fitness in RifR. We sampled actively growing bacterial cultures of each of the four strains, extracting total RNA and protein to be profiled using RNA sequencing (RNAseq) and sequential window acquisition of all theoretical mass spectra (SWATH-MS), respectively (see Figure 1B). In total, we were able to obtain RNA transcript counts for all present regions of the *Mtb* genome and reliably quantify 2,886 proteins across our samples (Figure S1). We used differential expression analysis to test our hypothesis that the compensatory mutation RpoC Leu516Pro had the net effect of reversing, at least partially, the expression changes brought about by the rifampicin resistance mutation RpoB Ser450Leu. We named this trend a “signature of compensation” – see Figure 2A and we derived it by identifying genes that are uniquely differentially expressed in RifR compared to the other three strains in our dataset. To maximise the probability of identifying the signature of compensation, we chose an inclusive definition of differential expression: a p-value of less than 0.05 after adjusting for multiple testing (see Methods). In keeping with our inclusive approach, we also deliberately did not use an effect size threshold (e.g. minimum log-fold change).

**Figure 2:**
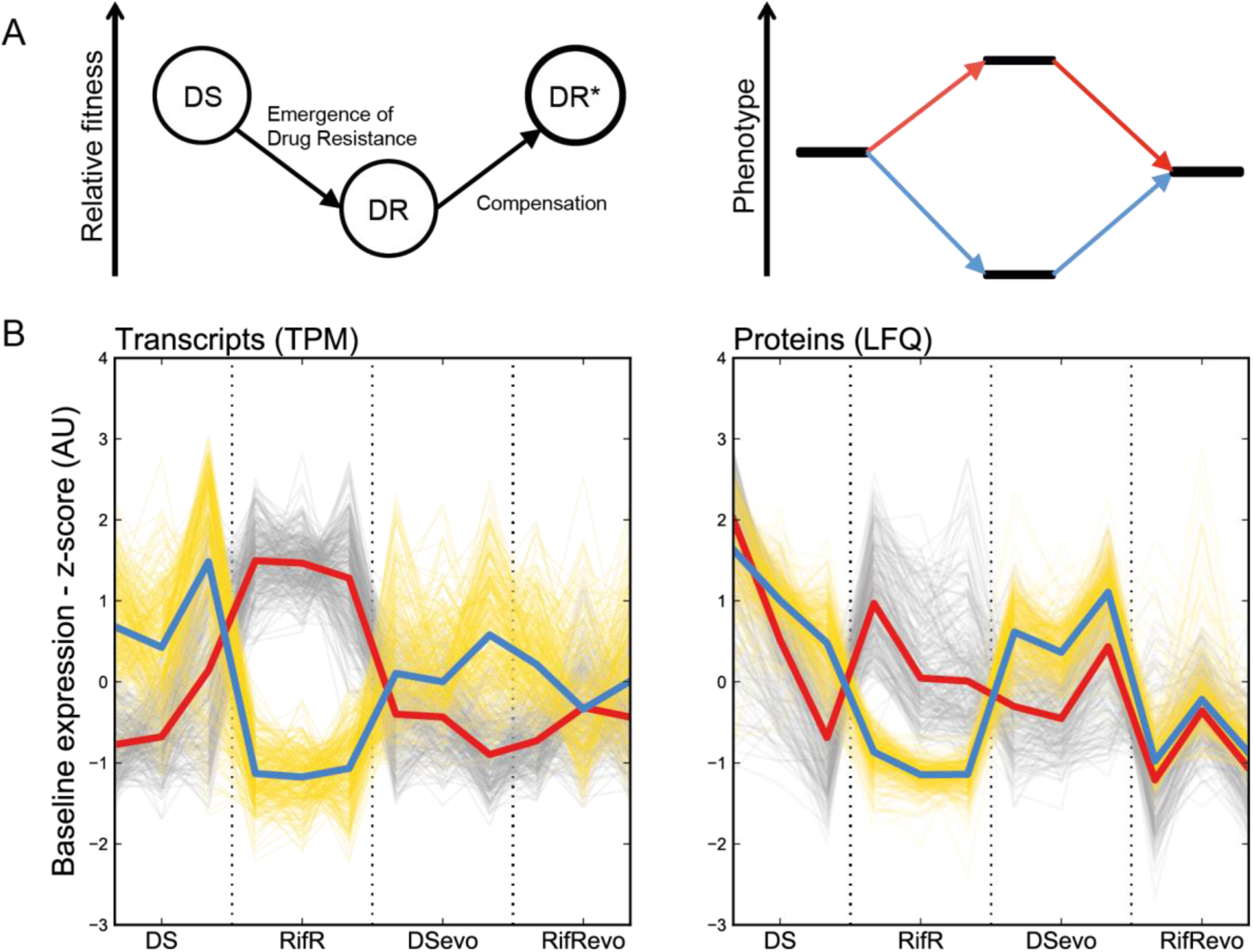
Signature of compensation. **A.** The relative fitness of drug resistant strains (DR) is expected to be lower than wild type (DS) at first, but then is expected to increase due to compensatory evolution. The phenotypic equivalent of this trend is illustrated as an increase/decrease in a measurable trait upon the emergence of resistance that is then returned to its previous level through compensation. We refer to this dynamic as the “Signature of Compensation”. **B.** Plot of transcript counts per million bases (TPM) and label free quantifications (LFQ) of cellular proteins for genes whose expression is perturbed by the Ser450Leu mutation in RpoB and returned to wild type in the presence of the compensating Leu516Pro mutation. All results were standardized across measurements for a single gene to allow the comparison between strains. Grey traces show genes that are significantly more highly expressed in RifR, yellow traces show genes that were significantly less highly expressed in RifR. The red and blue bold lines show the median of the sample for more and less highly expressed proteins, respectively. Data of three independent biological replicates for each strain are shown.

Using these criteria, we identified 536 transcripts that could be involved in the cost of resistance. 289 transcripts were less abundant and 247 were more abundant in RifR compared to the other samples. Similarly, 536 proteins showed a significant signature of compensation: 260 proteins were more and 276 were less-abundant in RifR (see Figure 2B). Gene set enrichment analysis of the transcriptomic and proteomic data pointed to iron homeostasis being significantly affected. Specifically, it indicated a higher expression, in RifR, of genes that are repressed by the iron-dependent regulator (IdeR, Rv2711) in iron replete conditions. Among them, there was a significant enrichment of genes involved in polyketide and non-ribosomal peptide synthesis, which include the biosynthetic machinery for the sole *Mtb* siderophore: mycobactin (see Figure S2-4). These changes suggested that RifR faced a shortage of iron in our experimental conditions.

The availability of iron is an essential requirement for *Mtb* growth, both in culture and during infection, and iron acquisition systems are therefore key virulence factors (Jones et al., 2014; Reddy et al., 2013; Wells et al., 2013). Hence, an increased requirement for iron could manifest itself as a loss of relative fitness. The fact that RpoB Ser450Leu led to a modification of the expression of genes involved in iron homeostasis and that RpoC Leu516Pro reversed the effect provides a compelling alternative mechanism underpinning the apparent fitness cost of rifampicin resistance. If the disruption of iron homeostasis drives fitness cost, we would expect that iron supplementation should mitigate the relative cost of RpoB Ser450Leu. Furthermore, based on the expression profile, we expected that RifR should produce more mycobactin at baseline than DS, potentially influencing the overall growth rate of the mutant.

We addressed the first hypothesis by comparing growth rates of RifR and DS in the presence or absence of 10 µM hemin – an additional source of iron that is by itself sufficient to support the growth of a mutant defective in mycobactin biosynthesis. Importantly, hemin and mycobactin provide two separate routes of iron uptake, which allows us to side-step issues that might emerge from deficient iron transport(Jones et al., 2014). The presence of hemin did not change the cost of RifR, which we calculated to be 18.6% in the absence and 20.9% in the presence of hemin for this experiment (Mixed effect linear model, p = 0.737). Similarly, hemin did not impact the growth rate of DS (−4.7%, CI^95%^: -16.3 – 2.3%, p = 0.128). In summary, iron did not appear to limit the growth of RifR under normal conditions.

Next, we addressed the production of mycobactin. We prepared whole cell extracts from DS and RifR grown in both, normal medium and medium supplemented with 10 µM hemin. We found that on average RifR produced more mycobactin than DS, corroborating the physiological relevance of the increased baseline expression of mycobactin biosynthesis genes. We also observed a slight decrease in the production of mycobactin in bacteria grown in the hemin-supplemented medium, pointing to a modification of the expression of mycobactin biosynthesis cluster in response to iron (See Figure 3). Given that the growth rate was not affected by the presence of hemin, these findings suggest that mycobactin itself does not modulate the growth rate of the mutant. It is therefore possible that the higher expression of the biosynthetic cluster itself might impart a fitness cost.

**Figure 3:**
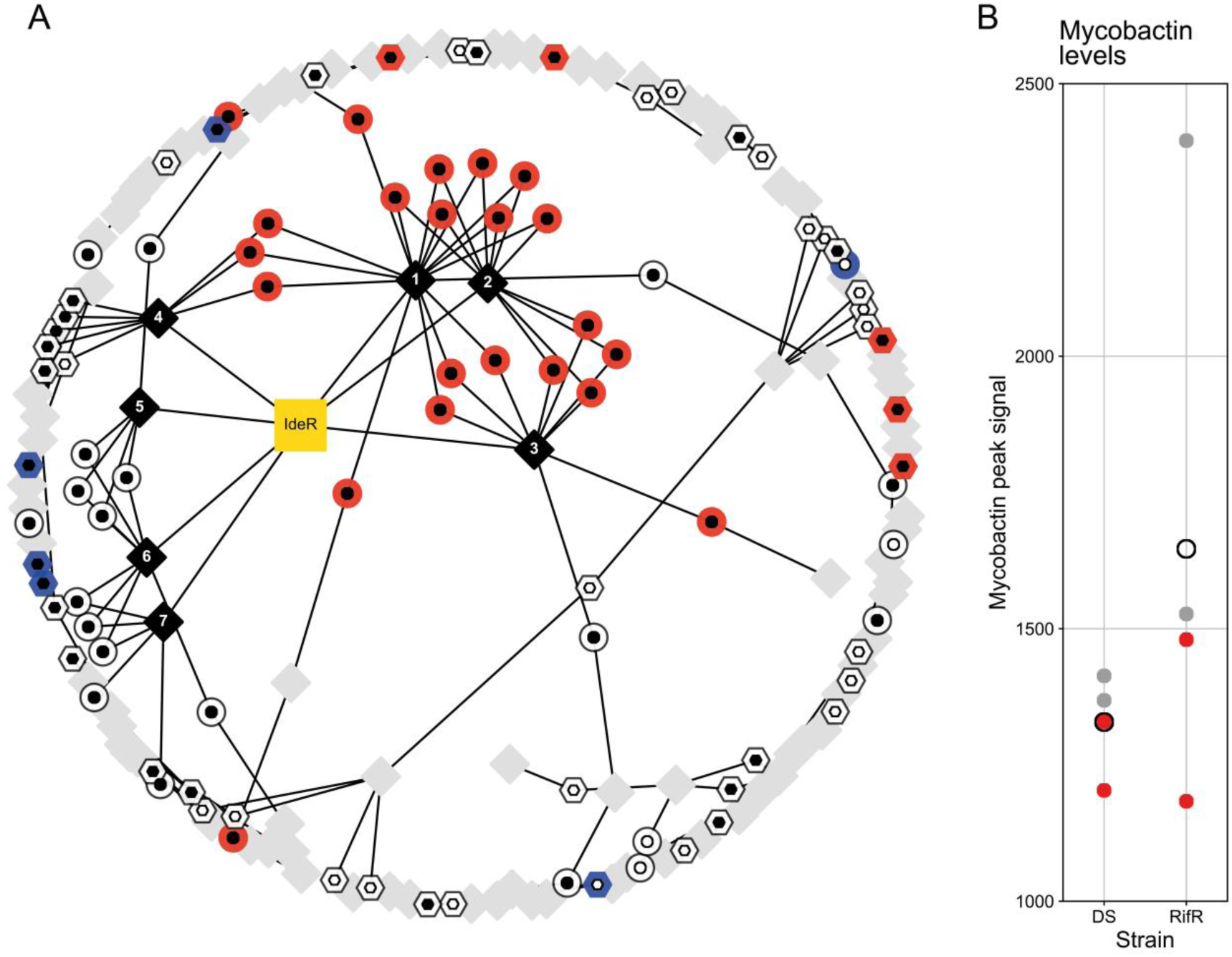
RifR has a higher baseline level of mycobactin biosynthesis than DS. **A**. Subset of the gene regulatory network (Peterson et al., 2014) containing iron responsive genes. Circles represent IdeR-regulated genes that are either induced (black inner circle) or repressed (white inner circle) in low iron conditions. Hexagons represent IdeR-independent iron responsive genes that are induced (white inner hexagons) or repressed (black inner hexagons) in low iron conditions. We used blue and red to indicate significantly lower or higher RNA expression in RifR, respectively – (n=12, see Methods for further details). Diamonds represent transcriptional modules as defined by Petersen et al, black diamonds indicate modules that contain at least 3 IdeR-responsive genes. Edges connect gene nodes with the module nodes they belong to. Labels 1-7 refer to Module 502 (1), Module 525 (2), Module 267 (3), Module 446 (4), Module 231 (5), Module 086 (6) and Module 295 (7) from the original publication. **B.** Relative mycobactin levels expressed as maximum peak heights for DS and RifR in normal medium (grey dots) and iron-supplemented medium (10µM hemin, red dots). Each filled circle represents the quantification of an independent biological replicate. Unfilled circles represent the mean of the observations.

Interestingly, while significantly enriched, only half of the genes reported to be repressed by IdeR (Rodriguez et al., 2002) in iron-replete conditions were part of the signature of compensation (22 out of 40 genes). This prompted us to take a closer look at the IdeR regulon and its regulation. We took advantage of recent studies modelling the global gene regulation in *Mtb* (Minch et al., 2015; Peterson et al., 2014; Rustad et al., 2014). We reconstructed the genome-wide gene regulatory network and extracted the immediate neighbours of IdeR- and iron-responsive genes(Peterson et al., 2014). There were 7 expression modules that contained at least 3 genes that are part of the IdeR regulon (Figure 3, black diamonds). Together, these modules covered 82.5% of all the IdeR-repressed genes, and with the exception of Module 4 (Figure 3), none of the modules included IdeR-independent iron-responsive genes. All the genes that we identified as candidates for compensation belonged to Modules 1-4, while none of the genes included in the other modules were found to be differentially expressed in RifR. A key difference among modules was that IdeR-regulated genes represented more than half of all the genes in modules affected by compensation but fewer than half in those that were not part of the “signature of compensation”. Mapping proteomic data onto the same expression network produced similar results (see Figure S5). Interestingly, few of the IdeR-independent iron-responsive genes were part of the signature of compensation. This pattern implies a modulation of the canonical function of IdeR, either through regulatory inputs from other transcription factors, or some other mechanism.

These results supported our hypothesis that mutations in *rpoB* impart changes to the baseline expression profile of *Mtb* that could be reversed in the presence of a compensatory mutation in *rpoC*. Combining the expression data with our findings that iron supplementation and mycobactin levels did not affect RifR growth rates, we concluded that the transcriptional changes were not driven by the demand for iron. Instead, these changes might be a reflection of a dysfunction of RNAP – e.g. differences in promoter specificity or modified interaction with IdeR, whose downstream consequences may impose a fitness effect. For example, as the mycobactin biosynthesis cluster comprises several large proteins, their excessive production could represent a drain on the cell’s resources. If true, we would expect such effects to be universal across all *Mtb* strains carrying this *rpoB* mutation.

### The impact of RpoB Ser450Leu is shaped by epistasis

We wanted to test the hypothesis that higher expression of the mycobactin biosynthetic cluster is a general feature of rifampicin resistance in *Mtb* and therefore the underlying cause of its fitness cost. To do so, we generated RpoB Ser450Leu mutants in five genetically diverse clinical isolates belonging to two different *Mtb* lineages and profiled them. Globally, *Mtb* can be grouped into seven distinct genetic lineages each with a specific geographic distribution (Gagneux, 2018). *Mtb* lineages can differ in their interaction with the human host, the dynamics of disease progression, and also in their apparent propensity to acquire drug resistance (Coscolla and Gagneux, 2014; Ford et al., 2013). We chose strains belonging to Lineage 1 and 2, because of their large phylogenetic separation (see Figure S6) and more importantly, because drug resistance is often associated with Lineage 2 and relatively rare in Lineage 1 (Borrell and Gagneux, 2009). We expected that the comparison of the transcriptome and proteome between the Ser450Leu mutants and their cognate wild type ancestor would allow us to identify general patterns of fitness cost linked to this mutation.

It is important to note that this comparison did not include any compensated strains, i.e. strains carrying mutations in the BBDP domain. We were therefore unable to focus our analysis exclusively on genes whose expression was corrected by the presence of an *rpoC* mutation. Nonetheless, direct comparison of RifR and DS is virtually indistinguishable from the signature of compensation when considering IdeR-regulated genes and therefore serves as a reasonable proxy for our analyses (see Figure S5).

We started by measuring the growth characteristics of the wild type isolates and the relative cost of the RpoB Ser450Leu mutation in the different strain backgrounds. The generation time varied from 22.7 h (^95%^CI: 20.8 – 25.0 h) to 31.0 h (^95%^CI: 29.3 – 35.1 h). The relative fitness cost of the RpoB Ser450Leu mutation differed as well, from a modest 2 % (mixed effect linear regression, p = 0.71) to a pronounced 27 % (mixed effect linear regression, p = 5.6 × 10^-6^).

We obtained the expression profiles for each strain to check whether the pattern we identified for IdeR-repressed genes was a universal phenotype for RpoB Ser450Leu mutants. Analysing the transcriptomic data by performing a single comparison across the five strain pairs, we found that only 17.5% (7/40 genes) of the IdeR-repressed genes were significantly differentially expressed. A single gene belonging to the mycobactin biosynthesis cluster was included in that number. Proteomic analysis revealed a similar result – 17.1% (6/35 detected proteins) were found to be significantly differentially expressed across all strains, none of which belonged to the mycobactin biosynthesis cluster. None of the iron-homeostasis gene sets highlighted in the “signature of compensation” were significantly differentially expressed across all strains. Since these findings were contrary to our expectations, we stratified the analysis and mapped the differential expression results for each strain onto the IdeR- and iron-responsive gene network we collated earlier. These results echoed our combined analysis: the signature of compensation was not universal across the tested strains. N0155, which corresponds to “DS”, is the only strain to show a transcriptional profile consistent with the signature of compensation (see Figure 4A). Proteomic data corroborated this finding (see Figure S7). It is important to note that these data represent an independent replication of the experiments, from which we derived the signature of compensation, showing that our original results are robust and reproducible. However, the absence of a coherent IdeR-responsive phenotype was clear evidence of epistasis and raised a broader question: are there any commonalities in the phenotypic manifestation of the RpoB Ser450Leu mutation among our set of strains?

**Figure 4:**
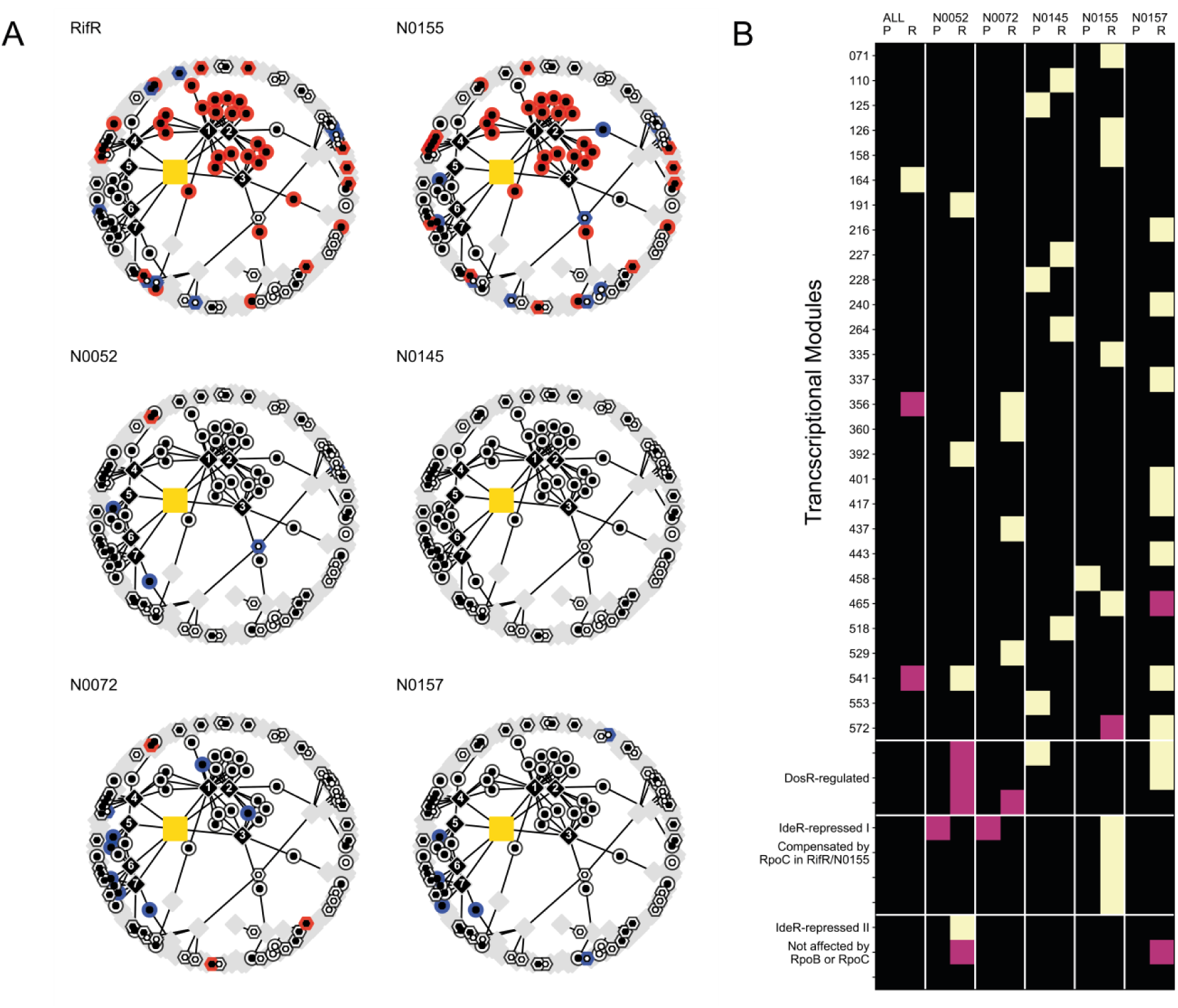
The prominent role of mycobactin biosynthesis in the signature of compensation is not universal. **A.** Iron-responsive subset the of gene regulatory network, as shown in Figure 3, coloured based on transcriptional differential expression data from pairwise comparison of genetically distinct rifampicin-susceptible clinical isolates and their cognate RpoB Ser450Leu mutants. RifR and N0155 refer to an independent sampling of the same strain pairs. See Figure S7 for the proteome counterpart of this plot. For RifR, N0072 and N0157 the plot is based on the comparison of three drug susceptible and three rifampicin resistant samples. For N0155, N0145 and N0052 we used two samples of each. **B.** Representation of the enrichment of significantly differentially expressed genes within individual transcriptional modules, as defined elsewhere (Peterson et al., 2014). The columns alternate proteomic (P) and transcriptomic data (R). “ALL” refers to the global differential expression analysis of all rifampicin-susceptible against all rifampicin-resistant strains. The remaining column annotations refer to individual pair-wise comparisons in different genetic backgrounds. Black squares represent no significant enrichment, mauve squares and yellow squares show enrichment at 0.01<p<0.05 and p<0.01 using a Fisher’s exact test. These p-values are not adjusted for multiple testing. Modules covering the DosR-regulon and IdeR-iron repressed regulon are highlighted separately.

To address this question we sought to identify expression modules (Peterson et al., 2014) whose membership was well represented among significantly differentially expressed genes in at least one pair-wise comparison between a rifampicin-resistant strain and its cognate drug-susceptible ancestor (see Methods for details). Using transcriptomic and proteomic data, we identified 33 expression modules that fitted our criterion (see Figure 4B). There was virtually no consensus across the strains in the transcriptional or translational response to the *rpoB* mutation. The only case where we observed partial agreement across genetic backgrounds concerned some of the modules controlled by the hypoxia-responsive regulator DosR(Park et al., 2003). As with modules containing IdeR iron-repressed genes, we observed only partial regulon induction for DosR. Specific modules were clearly involved in the expression changes (either protein or transcript) in each background, but the impact of these was strain-specific. A complementary manifestation of this phenomenon comes from the global comparison of all rifampicin-resistant strains against all wild type strains, which highlighted a single module as enriched for significantly differentially expressed genes. Comparing the distribution of the effect sizes, as measured by the per-gene fold-changes in expression in the combined analysis and the pairwise comparisons for each strain, we saw a marked muting of the magnitude of differential expression in the former (see Figure S8). This was likely due to the averaging effect of the combined analysis suppressing the contribution of the differential expression from individual strains. The magnitude of the expression change in pairwise comparisons was comparable across strains.

Overall, we were able to identify a wealth of gene expression changes in our samples: as many as 958 transcripts and 1914 proteins were observed to be differentially expressed in at least one comparison across our samples. On the level of individual genes, the transcriptome and to lesser extent the proteome of each strain were perturbed in their own private way (see Supplementary Figures 9&10), manifesting itself as the drug resistance iteration of the Anna Karenina principle (Zaneveld et al., 2017). Because the majority of those changes were specific to individual strains they were largely invisible if the comparison was made across all strain pairs. The fact that the same mutation can have such profoundly different outcomes depending on the genetic context in which it occurs, is clear evidence of epistasis, and shows that natural genetic variation can fundamentally impact the physiological consequences and therefore evolution of drug resistance. Importantly, the impact of resistance on the expression profile of any two strains was found to be independent of the genetic distance between them (see Figure S11).

So far, we showed that the RpoB Ser450Leu causes a considerable re-organization of baseline gene expression, that this perturbation can be reversed by a compensatory mutation in RpoC and that the specific phenotypic manifestation was dependent on mutations that occurred more recently than those defining individual lineages. These findings were consistent with our observation that the same mutation imposed a different fitness cost to different strains. We therefore sought to find correlates of the varying fitness costs.

### Deviation from baseline expression correlates with the cost of rifampicin resistance

Pleiotropic phenotypes of the kind described above are not normally addressed, however we wanted to explore whether the extent of the expression perturbations correlated with the varying fitness costs of Ser450Leu we observed in different genetic backgrounds. We reasoned that the cumulative impact on expression disruption, rather than the dysregulation of individual genes, would provide a conduit for a loss of fitness.

In the first instance, we considered the correlation between the fitness cost of the *rpoB* mutation and the overall expression distance between the mutant and its cognate wild type strain (See Figure S12). Through this approach, we were able to detect a relationship between cost and expression differences for the expressed proteins (R^2^= 0.83, p = 0.031, ordinary least squares linear regression) but not RNA (R^2^= 0.39, p = 0.258, ordinary least squares linear regression). Given that the correlation was stronger in the proteome compartment, and that the proteome compartment seemed more affected by resistance, we elaborated on our observation by incorporating a measure of physiological cost for each protein. We used two different metrics for cost. In the simpler case we used the molecular weight of amino acids as proxy for the resource investment necessary to generate each protein (Seligmann, 2003). We also used estimates of ATP cost for each amino acid in *E. coli* as a way to approximate the level of energy investment a bacterial cell makes when synthesising its proteome (Akashi and Gojobori, 2002). Both metrics showed that drug resistance imposes an additional physiological cost to the baseline proteome (Molecular Weight: Mann-Whitney U-test, p = 8.26 × 10^-4^, ATP equivalents: Mann-Whitney U-test, p = 4.50 × 10^-4^, see Figure S13). Furthermore, this cost was negatively correlated with the relative fitness of the RpoB Ser450Leu mutation in a given strain background (ϱ_s_ = -0.90, p = 0.04) – the greater the deviation from the resource investment of the ancestral proteome, the larger the cost of the mutation (see Figure 5A). Growth rate and gene expression are not independent from each other. To test the possibility that the observed correlation may be an artefact of our analysis, we took advantage of the natural variation in growth rates of different drug-susceptible clinical isolates in our medium and compared them to the relative costs of expression (See Figure S14). We performed a pairwise comparison across all the tested strains and observed no statistically significant correlation between the differences in the investment into the proteome and the difference in growth rates (ϱ_s_ = 0.34, p = 0.33). The differences in the allocation of resources into the protein compartment of different bacterial strains were therefore not the main determinant of variation in their respective generation times.

**Figure 5:**
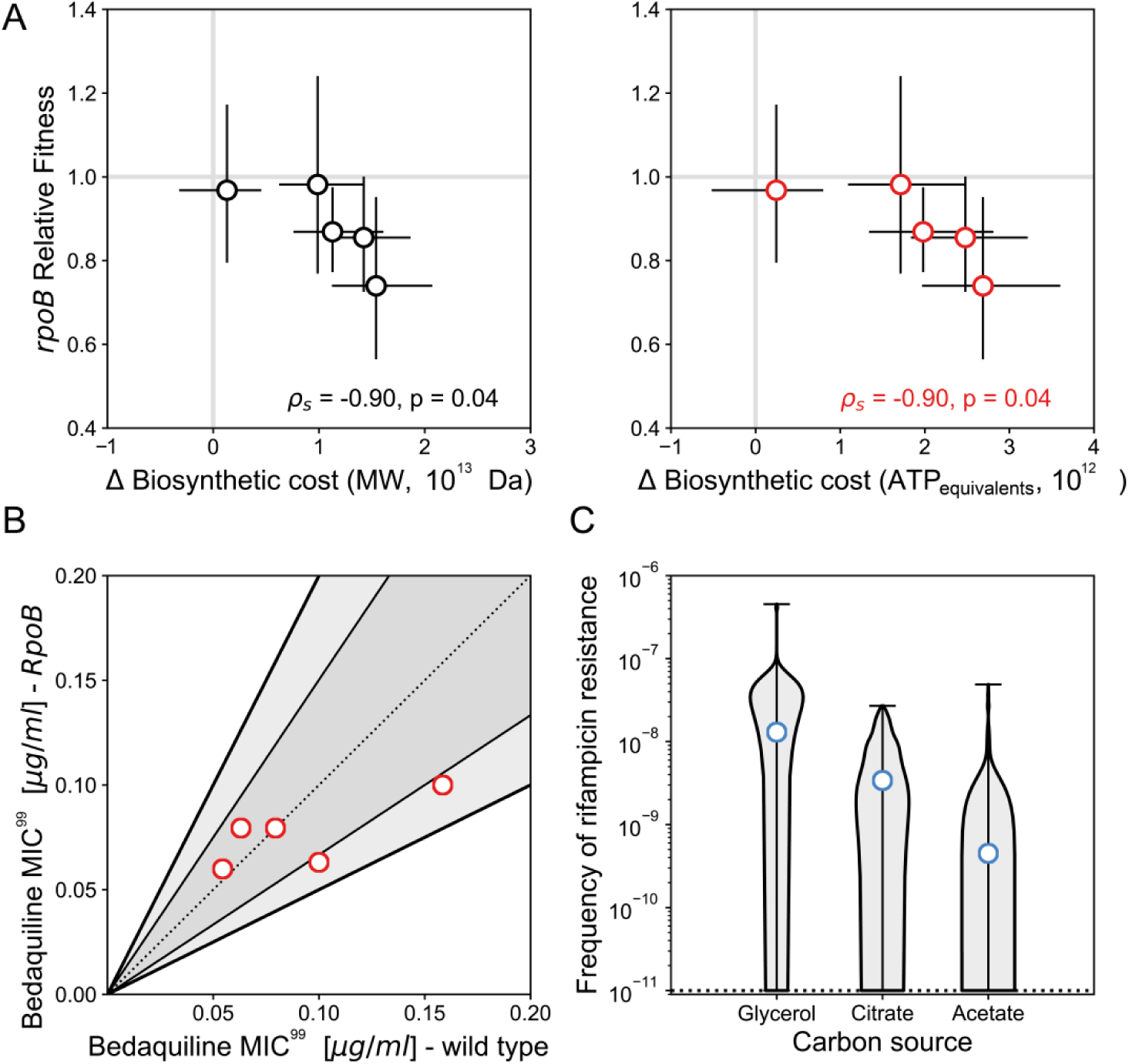
The fitness cost of RpoB Ser450Leu correlates with increased resource requirements. **A.** The relative fitness of Ser450Leu RpoB mutants estimated from growth rate data is negatively correlated with the magnitude of the deviation from the resources allocated to the wild type proteome. ϱs – Spearman correlation. Error bars indicate the 95% confidence interval for the data. Each cost determination was obtained from a minimum of four independent cultures per strain. Protein costs were derived from a total of 24 proteomic samples. **B.** Comparison of minimum inhibitory concentrations (MIC) of bedaquiline in clinical isolates and their cognate RpoB mutants. Dotted line shows parity, darker shading includes 50% or lower difference in MIC and the lighter shading spans up to 2-fold change in MIC. Each dot represents the mean of three independent measurements. **C.** The frequency of rifampicin resistance as measured in the model organism *Mycobacterium smegmatis* with the Luria-Delbrück fluctuation assay in media containing different carbon sources. The blue dot corresponds to the Ma-Sandri-Sarkar maximum likelihood estimate for the frequency of rifampicin resistance. The shaded area shows the density distribution of the number of resistant colonies per scored culture. The dotted line indicates the limit of detection. Each estimate is based on 120 independent cultures for glycerol and citrate and 150 independent cultures for acetate.

Taken together, our results seemed to suggest that the ultimate manifestation of the disruption of wild type baseline gene expression by RpoB Ser450Leu was a net increase in the biosynthetic input required to maintain the steady state proteome: the greater the cost of the disruption, the greater the slowing down of growth in a given strain background. We propose this as the “Burden of Expression” hypothesis of the fitness cost of rifampicin resistance.

### Carbon allocation rather than ATP availability modulates cost of resistance

An implication of the “Burden of expression” hypothesis is the possibility of suppressing the emergence of rifampicin-resistance in mycobacteria by maximising the additional biosynthetic cost imposed by the deviation from the baseline expression. We tested two types of conditions that may impose such a stress: inhibition of ATP synthesis and variation of carbon-source quality. The first would disrupt the ability to generate energy through catabolic processes, while the second would place more emphasis on the anabolic aspects of bacterial growth. In the first instance, we tested the susceptibility to bedaquiline, an ATP synthase inhibitor that leads to a decrease in intracellular ATP levels in *Mtb* (Andries et al., 2005). Given the higher baseline cost of their proteome, we expected that RpoB Ser450Leu mutants should show an increased susceptibility to bedaquiline commensurate with their relative loss of fitness. We did not observe any correlation between bedaquiline susceptibility and the cost of the RpoB Ser450Leu mutation (see Figure 5B).

Next, we explored varying carbon source quality, expecting substrates that force the bacterial cell to rely more heavily on anabolic processes to serve as amplifiers for the perceived cost of rifampicin resistance. A related phenotype has been reported before for RpoB Ser450Leu(Song et al., 2014). We chose the Luria-Delbrück fluctuation assay as an unbiased readout for the overall increase in the cost of rifampicin-resistance, because its frequency of resistance estimate contains a signal for the ability of drug resistant bacteria to propagate within the population prior to antibiotic exposure(Ycart, 2013). The global increase in the cost of RpoB mutations would therefore manifest itself as an apparent decrease in the frequency of resistance, as the population size of pre-existing RpoB mutants would be smaller due to limited expansion post-emergence. We chose glycerol, citrate and acetate to test our hypothesis in the soil organism *Mycobacterium smegmatis*, whose patterns of rifampicin resistance mirror those of *Mtb* (Borrell et al., 2013). As expected, these three carbon sources supported different growth rates with measured generation times of the wild type being 3.24 h (^95%^CI: 3.23 – 3.25 h), 6.17 h (^95%^CI: 6.09 – 6.25 h) and 17.62 h (^95%^CI: 17.61 – 17.62 h), respectively. We then determined the frequency of rifampicin resistance for bacteria grown on each carbon source using the Luria-Delbrück fluctuation assay. We found a striking correlation between carbon source and the calculated frequency of resistance, with bacteria grown in glycerol giving rise to rifampicin-resistant bacteria at a rate of 1.3 × 10^-8^(^95%^CI: 1.2 × 10^-8^– 1.5 × 10^-8^), those grown in citrate at a rate of 3.4 × 10^-9^(^95%^CI: 2.9 × 10^-9^– 4.0 × 10^-9^) and acetate-cultured bacteria at a rate of 4.5 × 10^-10^(^95%^CI: 3.4 × 10^-10^– 5.6 × 10^-10^) – see Figure 5C. This trend was remarkable, because it showed that changing only the carbon source, keeping all other variables constant, could lead to a 28-fold change in the frequency of resistance.

The disparity in outcomes between the two experimental approaches suggests that the availability of catabolic energy does not disproportionately influence the ability of RpoB mutants to survive. However, the impact of carbon source on the frequency of rifampicin-resistant bacteria within a population clearly suggests that carbon allocation might be an important driver of the fitness cost of rifampicin resistance.

## Discussion

We normally expect that form follows function in bacteria: expression differences should reflect variations in physiological states. Indeed, we show that RpoB Ser450Leu imparted a measurable physiological perturbation in addition to conferring rifampicin resistance. Consistent with the suggested role of compensatory mutation (Comas et al., 2012), we confirmed that in one strain, RpoC Leu516Pro reduced both, the apparent fitness cost of rifampicin resistance and the magnitude of the expression changes arising from it. However, we also showed that the nature of the perturbation was not consistent across different genetic backgrounds. Instead, we observed a strain-specific response to the RpoB mutation, both in terms of the relative impact on growth and the rearrangement of gene expression. We further observed that the magnitude of the fitness cost that RpoB Ser450Leu imposes on a strain was related to the overall increase in the resources allocated to the proteome. Based on these observations, we proposed the “Burden of expression” hypothesis, with which we posited that in *Mtb*, the cost of rifampicin resistance was mediated by the metabolic burden imposed by the modified baseline protein expression of resistant strains. Elaborating on this hypothesis we demonstrated that interfering with anabolic processes could suppress the emergence of rifampicin resistance in the related organism *M. smegmatis*.

The “Burden of expression” hypothesis stems from experimental data with clear caveats. First, we started our analyses assuming that ribosomal biosynthesis is unlikely to play a key role in the cost of rifampicin resistance in *Mtb* and that therefore expression data were a better window into the modified physiology. Our data seem to support the validity of this assumption: ribosomal proteins represented only 5.5%, on average, of the total protein biomass in our experiments. This proportion was marginally higher in RpoB mutants, and it seemed to increase with increasing generation time (see Figure S15). These trends were more consistent with a cost imposed by the metabolic burden of making ribosomes. Second, some of our key conclusions are based on a relatively small number of strains. Nonetheless, to the best of our knowledge, this sample set represents the most comprehensive and best curated account of rifampicin resistance-induced global expression changes in *Mtb* to date, covering both: evolutionary dynamics and phylogenetic diversity. We were also able to show that patterns of expression detected in the DS-RifR comparison were robust when the same strain pair was sampled again (see Figure 4 and Figure S7). Importantly, key inferences that led us to propose the hypothesis came from SWATH-MS proteomic data drawn from the five different strain backgrounds. These data showed a clear clustering of biological replicates (see Figure S16), with the exception of N0145 for which we were also unable to detect a significant cost for the Ser450Leu mutation or any significant changes to the expression. Third, we assumed that label free quantification (LFQ) using the “best flyer peptide” or TopN approach, which reflects the proportional abundance of individual proteins within our samples (Schubert et al., 2015), can be used to draw conclusions about the resource investment of the cell and can be extended to the growth rate of bacteria. It is possible that the roles are reversed and the growth rate of bacteria in fact determines the protein complement being expressed (Beste et al., 2007). We addressed this possibility by performing a comparison of proteome investment and growth rate for wild type strains only. If the growth rate of *Mtb* did indeed determine the protein complement of cells across genetic distances on an evolutionary timescale, we would expect a strong correlation between differences in proteome and differences in growth rates between any two strains. This was however not the case (see Figure S13). Finally, we also assumed that the proteome plays a central role in imposing a limit to the growth rate of an *Mtb* cell. There are other components that require considerable investment in carbon: in the case of *Mtb* both lipids and cell wall may act as a sink for resources limiting growth as they can account for over half of the dry mass of actively growing cells(Beste et al., 2005). Lipidomic analysis of RpoB mutants in *Mtb* pointed to differences in mycobactin biosynthesis as one of the biggest discrepancies between rifampicin-resistant mutants and their susceptible ancestors (Lahiri et al., 2016). While echoing a key observation from our quest for determining the cost of resistance, we saw no evidence that mycobactin biosynthesis itself changes the rate of bacterial growth. The virulence-associated phthiocerol dimycocerosates (PDIM) have also been implicated in the cost of rifampicin resistance (Bisson et al., 2012), as have other changes in lipid composition (du Preez and Loots du, 2012). The full exploration of the role of lipids in the physiology of rifampicin-resistant *Mtb* is beyond the scope of this study, but it would provide an interesting new and complementary avenue to pursue.

Keeping these considerations in mind, there are two striking features to emerge from our results. The first is the pervasive epistasis modulating the impact of RpoB Ser450Leu: the same mutation has markedly different effects on the physiology of different *Mtb* strains. The second is the apparent mechanism through which modulation of gene expression is propagated across the levels of bacterial physiology. Modification in RNAP function seems to have pleiotropic effects that transcend the disruption of any single group of genes, and impart a perturbation that appears to affect bacterial resource allocation.

One question that remains open is what sits at the heart of the disparity in phenotypes? The sequence of RNAP is effectively the same in all strains (Borrell and Trauner, 2017); and by extension so are the biochemical changes that arise from resistance (Stefan et al., 2018). We envisage that part of the answer lays in differences in underlying robustness: a strain’s capacity to buffer perturbation. Furthermore, we can consider this a window into the evolutionary adaptation of each strain and a sign of how different their physiologies really are. The amalgamation of mutational differences that effectively makes up a strain genetic background weaves a baseline phenotype that allows different *Mtb* strains to be successful pathogens despite differences in their underlying physiology: i.e. there are several successful approaches to solving the same problem. These differences are unmasked by the presence of a mutation that sits at the core of gene expression and reveals idiosyncratic transcriptional responses to rifampicin resistance that are poorly conserved across genetic distances. This observation has the implication that, beyond the described mutations in BBDP, which seem to alleviate some of the biochemical and gene expression effects of rifampicin resistance more generally, further investigation of positive selection of compensation of resistance-related traits should be performed in genetically related strains as they could vary considerably when comparing phylogenetically distant strains (Farhat et al., 2013; Zhang et al., 2013).

The strain-specific nature of resistance-related expression perturbations can be used to provide a credible link to disparate growth rate modulation. Our suggestion that proteome composition influences growth rate is not without precedent. This connection has been made before (Scott et al., 2010), and resulted in the formulation of a collection of “growth laws” that linked growth rates to the partitioning of the limited proteome between ribosomes and other proteins carrying out the rest of the cellular functions. Growth on different carbon sources impacted this balance, with “poorer” ones requiring a greater investment into the functional proteome, presumably because of the need for anabolic reactions increased the reliance on biosynthetic enzymes. A similar relationship has been observed in a wide range of microbial species (Karpinets et al., 2006). An elaboration of these growth relationships also led to the conclusion that the efficiency of proteome allocation can impact growth rates and cell physiology (Basan et al., 2015). Our finding that the increase in the relative cost of the proteome brought about by the gain of a mutation correlates with the relative fitness of that mutation is consistent with these reports, as is our observation that anabolic processes may play a mechanistic role in setting the cost of a mutation.

The observed differential cost of rifampicin resistance across *Mtb* strains, provides a lens through which we can better understand the emergence of drug resistance in clinical TB. However, it also indicates a new avenue to pursue in the fight against rifampicin resistant *Mtb* and perhaps uncover a new paradigm for chemotherapeutic intervention. Agents that impart a considerable shock to the expression equilibrium of bacteria could exhibit potent activity against rifampicin resistant strains due to collateral sensitivity. Furthermore, when given in combination with rifampicin, such agents may act to suppress the emergence of resistance; a valuable attribute for lengthening the shelf life of rifampicin.

## Methods

### Strains and culture conditions

We used four strains described by Comas *et al.*(Comas et al., 2010): namely the wild type, clinical isolate T85 (N0155, DS), a rifampicin resistant mutant of T85 carrying the Ser450Leu mutation (N1981, RifR), a derivative of T85 that was evolved by serial passage (200 generations) in the absence of rifampicin (N1588, DS^evo^) and an evolved derivative of the rifampicin resistant strains carrying an additional mutation in RpoC – Leu516Pro (N1589, RifR^evo^).

In addition to these strains we used four clinical isolates that are part of the recently compiled Reference set of *Mtb* clinical strains(Borrell et al., 2018) covering the genetic diversity of *Mtb*. Two strains belonging to Lineage 1 (N0072, N0157) and two to Lineage 2 (N0052, N0145). We plated each of these strains on 7H10 plates containing 5 μg/ml Rifampicin, and picked colonies of spontaneous mutants. We checked the rifampicin-resistance conferring mutations using Sanger sequencing of the amplified RRDR region (Forward primer: TCGGCGAGCTGATCCAAAACCA, Reverse primer: ACGTCCATGTAGTCCACCTCAG, product size: 601 bp), and kept a Ser450Leu derivative of each clinical strain (N2027, N2030, N2495 and N1888, respectively).

Bacteria were cultured in 1l bottles containing large glass beads to avoid clumping and 100 ml of media incubated at 37°C rotated continuously on a roller. Unless otherwise stated we used a modified 7H9 medium supplemented with 0.5% w/v pyruvate, 0.05% v/v tyloxapol, 0.2% w/v glucose, 0.5% bovine serum albumin (Fraction V, Roche) and 14.5 mM NaCl. Compared to the usual composition of 7H9 we omitted glycerol, tween 80, oleic acid and catalase from the medium. We added 10 μM Hemin (Sigma) when supplementing growth medium with iron. We followed growth by measuring optical density at 600 nm (OD_600_).

Fluctuation assay experiments were performed using *Mycobacterium smegmatis*, mc^2^155. *M. smegmatis* was grown either in 10 ml cultures within 50 ml Falcon conical tubes in a shaker incubator (37°C, 200 rpm), or as 200 μl aliquots within flat-bottomed 96-well plates at 37°C and shaken at 200 rpm. We followed growth by measuring OD_600_. We used unmodified 7H9 medium or medium where glycerol was replaced with citrate or acetate added at concentrations that matched the molarity of carbon.

### Fitness determination

*Mtb* fitness was determined by comparative growth rate estimation. We grew bacteria as described and followed their growth by measuring OD_600_ with a Ultrospec 10 (GE Lifesciences). We transformed the optical density measurements using logarithm base 2 and trimmed all early and late data points that deviated from the linear correlation expected for exponential growth. Next, we fitted a linear mixed effect regression model to the data. Fitness cost was calculated as the resistance imposed deviation from wild type growth dynamics.

For *M. smegmatis*, we determined the growth rates by culturing bacteria as described above. We monitored the increase in OD_600_ using a Tecan M200 Pro Nanoquant at 20 min intervals. The data were log2-transformed, trimmed to retain only the portion of data pertinent to exponential growth and used for fitting a mixed effect linear regression model to estimate growth parameters.

### Transcriptional analysis with RNAseq

We transferred a 40 ml aliquot of bacterial culture in mid-log phase (OD_600_ = 0.5 ± 0.1) into a 50ml Falcon conical tube containing 10 ml ice. We harvested the cells by centrifugation (3,000×*g*, 7 min, 4°C), re-suspended the pellet in 1 ml of RNApro solution (MP Biomedicals) and transferred the suspension to a Lysing matrix B tube (MP Biomedicals). We disrupted the bacterial cells using a FastPrep24 homogeniser (40s, intensity setting 6.0, MP Biomedicals). We clarified the lysate by centrifugation (12,000×*g*, 5 min, 4°C), transferred the supernatant to a clean tube and added chloroform. We separated the phases by centrifugation (12,000×*g*, 5 min, 4°C) and precipitated the nucleic acids from the aqueous phase by adding ethanol and incubating at - 20C overnight. We performed a second acid phenol extraction to enrich for RNA. We treated our samples with DNAse I Turbo (Ambion), and removed stable RNAs by using the RiboZero Gram Positive ribosomal RNA depletion kit (Epicentre). We prepared the sequencing libraries using the TruSeq stranded Total RNA kit (Illumina) and sequenced on a HiSeq2500 high output run (50 cycles, single end).

Illumina short reads were mapped to the *Mtb* H37Rv reference genome using BWA(Li and Durbin, 2010) (ver 0.7.13); the resulting mapping files were processed with samtools(Li et al., 2009) (ver 1.3.1). Per-feature read counts were performed using the Python module htseq-count(Anders et al., 2015) (ver 0.6.1p1) and Python (ver 2.7.11). We performed differential expression analysis using the R package DESeq2(Love et al., 2014) (ver 1.16.1) and R (ver 3.4.0). In the case of the identification of the signature of compensation we performed a comparison of RifR vs DS + DS^evo^+ RifR^evo^. For the follow-up experiments we performed two separate comparisons: (DR^N0072^ + DR^N0157^ + DR^N0052^ + DR^N0145^ + DR^N0155^) vs (DS^N0072^ + DS^N0157^ + DS^N0052^+ DS^N0145^+ DS^N0155^) as well as individual DR vs DS comparisons.

Gene set enrichment analysis was based on functional annotation from the Kyoto Encyclopaedia of Genes and Genomes and a custom collation of curated gene sets based on published reports. The overrepresentation analysis was based on Fisher’s exact as the discriminating test.

In addition we transformed per-feature counts into transcript counts per million bases (TPM). TPM for each feature for each sample were calculated using the following formula:

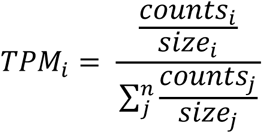

Where *counts*_*i*_ refers to the number of reads that map to a feature *i*, and *size*_*i*_ refers to the length (in bp) of feature *i*. This ratio was normalized by dividing by the sum of all the ratios across all the features.

### Proteomic analysis with SWATH-MS

We harvested 20 OD_600_ equivalents from mid-log phase (OD_600_ = 0.5 ± 0.1) bacterial cultures by centrifugation (3,000×*g*, 7 min, 4°C). We washed the bacterial pellet twice with phosphate buffered saline (PBS) to remove residues of tyloxapol. We re-suspended the bacterial pellet in 500 μl of protein lysis buffer (8M Urea, 0.1 M Ammonium bicarbonate, 0.1% RapiGest [Waters]) and transferred the suspension to a Lysing matrix B tube (MP Biomedicals). We disrupted the bacterial cells using a FastPrep24 homogeniser (40s, intensity setting 6.0, MP Biomedicals). We clarified the lysate by centrifugation (12,000×*g*, 5 min, 4°C), and sterilised the supernatant by passing it twice through a 0.22 μm syringe filters (Milipore).

Following protein extraction for each sample, we used trypsin to digest proteins into peptides and then desalted them using C_18_ columns (The Nest Group). The cleaned up peptides were re-suspended in MS buffer (2% v/v acentonitrile, 0.1% v/v formic acid). Finally, the RT-kit (Biognosis) containing 11 iRT retention time normalization peptides was spiked in to every sample.

We measured every sample in sequential window acquisition of all theoretical mass spectra (SWATH) mode, a data independent acquisition implementation, on a tripleTOF 5600 mass spectrometer (AB Sciex) coupled to a nano flow HPLC system with the gradient of one hour(Banaei-Esfahani et al., 2017). The raw files acquired through a 64 variable width window precursor isolation scheme were centroid normalized using Proteowizard msconvert. We used the *Mtb* spectral library described earlier(Schubert et al., 2013) to extract data using the OpenSWATH workflow(Reiter et al., 2011; Rost et al., 2014; Rost et al., 2016). The processed data were filtered by MAYU to 1% protein FDR(Reiter et al., 2009). R packages aLFQ and MSstats were used for protein quantification (Top3 peptides and top5 fragment ions(Schubert et al., 2015)) and differential expression analysis respectively(Choi et al., 2014; Rosenberger et al., 2014).

### Mycobactin determination

We harvested 5 OD_600_ equivalents from mid-log phase (OD_600_ = 0.5 ± 0.1) bacterial cultures by centrifugation (3,000×*g*, 7 min, 4°C). We washed the bacterial pellet three times with 15ml of cold, sterile 7H9 medium base devoid of additives (BD) to remove residues of tyloxapol. After washing we resuspended the pellets in 80 μl of cold, sterile 7H9 medium base and added 750 μl of 1:2 Chloroform:Methanol. We vortexed the samples for 5 minutes at top speed and added 750 μl of Chloroform. The samples were shaken for 1.5h at room temperature and clarified by centrifugation (16,000 × *g*, 10 min). We transferred the organic phase to a fresh tube, dried the samples in a speedvac and re-suspended each sample in 120 μl of 44:44:2 Acetonitrile:Methanol:H2O, (v:v:v).

Chromatographic separation and analysis by mass spectrometry was done using a 1200 series HPLC system with a Phenomenex Kinetex column (1.7 µl × 100 mm × 2.1 mm) with a SecurityGuard Ultra (Part No: AJ-9000) coupled to an Agilent Technologies 6550 Accurate-Mass Q-Tof. Solvent A: H_2_O, 10mM ammonium acetate; Solvent B: acetonitrile, 10mM ammonium acetate. 10 µl of extract were injected and the column (C18) was eluted at 1.125 ml/min. Initial conditions were 60% solvent B: 0-2 min, 95% B; 2-4 min, 60% B; 4-5 min at initial conditions. Spectra were collected in negative ion mode form 50 – 3200mz. Continuous infusion of calibrants (Agilent compounds HP-321, HP-921, HP-1821) ensured exact masses over the whole mass range.

We converted the raw data files to the mzML format using msConvert and processed them in R using the XCMS(Smith et al., 2006) (ver 3.0.2). We extracted targeted ion chromatograms with CAMERA(Kuhl et al., 2012) (ver 1.34.0).

### Transcriptional module analysis

The iron-responsive sub-graph of the global gene regulation network published by Peterson *et al.*(Peterson et al., 2014), was generated by using all expression modules and all iron-responsive genes as nodes, with edges connecting them representing module membership. All other gene nodes were discarded, keeping only the information pertinent to the number of genes present in each module (its degree). We focused explicitly on modules with at least 3 IdeR-dependent iron-responsive genes within them. Finally we marked significant differential expression of the gene nodes in every comparison.

For the purposes of contextualising the expressional profiling of RpoB Ser450Leu we selected a subset of expression modules as follows: first we collated all the genes that were differentially expressed in at least one genetic background as determined by pairwise comparisons. We then scored each expression module for enrichment of membership by differentially expressed genes using a binomial test. We retained all modules for which the test pointed to an excess of differentially regulated genes (p < 0.05). We constructed a new sub-graph of the global regulatory network using all enriched modules and their constituent genes irrespective of whether or not individual genes were significantly differentially expressed. Edges reflected module membership. We added expression information in the form of log-fold changes of abundance to each subgraph based on pairwise analyses.

### Calculation of genetic distance between clinical isolates

Genetic distance between strains was defined as the number of single nucleotide variants (SNV) that separate two strains. The numeric value of this parameter was extracted from the phylogeny published elsewhere(Borrell et al., 2018).

### Quantification of the relative impact of the *rpoB* mutation on gene expression in different clinical isolates

We define the dissimilarity in the expressional response to the presence of the *rpoB* mutation using three metrics: absolute number of shared significantly differentially expressed genes, the fraction of both the shared significantly differentially expressed genes and shared non-affected genes (hamming distance) and the Euclidean distance between ratios of TPM. The first is simply the number of shared genes that were found to be significantly affected by the presence of the *rpoB* mutation in two different genetic backgrounds. For the second we use the same input to calculate the hamming distance between the patterns of genes significantly affected by the mutation in *rpoB* in two different genetic backgrounds. In the third case we first calculate the TPM. We then calculate the mean TPM for each gene across the biological replicates as well as the ratio of mutant to wild type mean TPM for every gene. This gives us a vector containing 4000 ratios for each mutant-wild type pair. Finally we calculate the Euclidean distance between these vectors for the different genetic backgrounds. We plotted each of these metrics against genetic distance and calculated the spearman correlation and the coefficient of variance: standard deviation over mean multiplied by 100 (σ / μ × 100%).

### Quantification of the absolute impact of the *rpoB* mutation on gene expression of a clinical isolate

We used transcript counts per million bases (TPM) and label free quantification (LFQ) to generate an RNA vector and a protein vector containing all the available information for each measured sample. We then calculated all the possible DS – RifR pairwise Euclidean distances for the RNA and protein vectors within each genetic background. We used the mean and standard deviation for the dissimilarity estimates. We evaluated the correlation between the fitness cost of RpoB mutations and the expression distance using the R^2^-coefficient derived from ordinary least squares linear regression as well as the Spearman correlation. Arbitrary units expressing the dissimilarity were obtained by dividing the calculated distances by 500,000 or 10,000,000 for TPM and LFQ, respectively.

### Estimation of the biosynthetic cost of protein production

The calculation of biosynthetic cost was based on the molecular weight of amino acids (MW)(Seligmann, 2003) or on the estimate of *E. coli* ATP investment into individual amino acids derived by Akashi *et al.*(Akashi and Gojobori, 2002) using the following formulae:

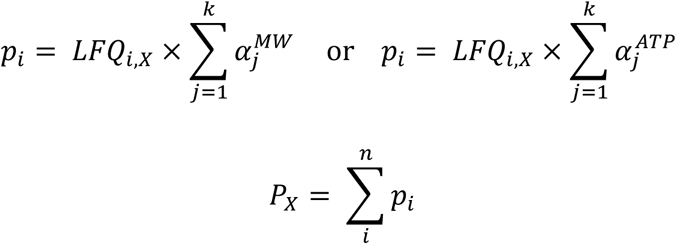

Where the cost of protein *i* (*p*_*i*_) was calculated as the sum of the cost for each constituent amino acid 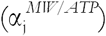 based either on its molecular weight (*MW*) or ATP investment (*ATP*) and adjusted by the proportional contribution of protein *i* to the total proteome of sample *X* (*LFQ*_*i,X*_). The overall cost of the proteome *P* for a sample *X* (*P*_*X*_) is expressed as the sum of the costs of individual proteins (*p*). The difference between the biosynthetic investments in the proteome of sample X when compared to sample Y was simply: *P*_*X*_ – *P*_*Y*_. We estimated the biosynthetic perturbation of RpoB Ser450Leu within a genetic background, by resampling sample-specific proteome costs for DS and RifR with replacement 100-times, and using the median as well as the 3^rd^and 98^th^quantiles to provide the 95% confidence interval. Finally, we quantify the correlation with the relative fitness of RpoB Ser450Leu by calculating the Spearman coefficient.

### Minimum inhibitory concentration determination

We used the microplate alamar blue assay(Franzblau et al., 1998) to determine the minimum inhibitory concentrations of bedaquiline in all drug susceptible and drug resistant strains used in our study. We tested bedaquiline using a two-fold dilution series spanning a concentration of 4 ng/ml – 1 µg/ml.

### Fluctuation Assay for determining the frequency of rifampicin resistance

We used the Luria-Delbrück fluctuation assay(Luria and Delbruck, 1943) to determine the frequency of rifampicin resistance in *Mycobacterium smegmatis*. Briefly, we inoculated 30 parallel cultures containing 10 ml of modified Middlebrook 7H9 medium containing either glycerol, citrate or acetate as the main carbon source with 5000 colony forming units of pre-adapted *M. smegmatis*. We grew the cultures to mid-log phase (OD_600_=0.5) at which point we chose three cultures at random for the determination of overall population size. We harvested the remaining bacteria by centrifugation 4000×*g* for 7 minutes, re-suspended the cellular pellet with 500 µl of fresh Middlebrook 7H9 medium and plated onto Middlebrook 7H10 solid media supplemented with 200 µg/ml Rifampicin. Plates were incubated at 37°C for 3-4 days and scored by counting the resulting resistant colonies. We determined the population-wide number of mutants (*m*) using an in house implementation of the Ma-Sandri-Sarkar maximum likelihood estimation(Sarkar et al., 1992), and adjusted it by the estimated population size to determine the frequency of resistance.

### Quantification and statistical analysis

Unless otherwise stated we preformed the analyses using Python 3.5.2 augmented with the following modules to provide additional functionality: Matplotlib (ver 2.0.0), Numpy (ver 1.12.1), Scipy (ver 0.19.0), Pandas (ver 0.20.1), statsmodels (ver 0.8.0), sklearn (ver 0.18.1), and netwrokX (ver 1.11). All the details pertaining to the statistical treatment of data can be found where results are described: either in the main text, figure legends or methods.

### Data and Software availability

All RNAseq data were deposited in the ArrayExpress repository of the European Bioinformatics Institute under the E-MTAB-7359 accession. The mass spectrometry proteomics data have been deposited to the ProteomeXchange Consortium via the PRIDE partner repository with the dataset identifier PXD011568. These data are pertinent to Figures 2-5 and all Supplementary Figures with the exception of Figure S6.

A record of data analysis pertinent to this paper will be made available at http://www.github.com/swissTPH/TBRU_cost_of_resistance/.

## Acknowledgements

Calculations were performed at sciCORE (http://scicore.unibas.ch/) scientific computing center at University of Basel, with support by the SIB - Swiss Institute of Bioinformatics. This work was supported by the SystemsX.ch project “TbX”, the National Institutes of Health project Omics4TB Disease Progression (U19 AI106761), Swiss National Science Foundation (grants 310030_166687, IZRJZ3_164171, IZLSZ3_170834 and CRSII5_177163) and the European Research Council (309540-EVODRTB). The authors would like to thank Uwe Sauer and Michael Zimmermann for their input during the early stages of the project. We would like to thank Janssen Pharmaceutica NV for their kind gift of bedaquiline.

## Author Contributions

AT, RA and SG designed the project and wrote the manuscript. AT, SMG and SB generated samples for expression profiling, mycobactin determination and measured the fitness of *Mtb* strains. JF obtained MICs for bedaquiline. SS measured the frequency of resistance in *M. smegmatis*. PW and MZ performed the sample acquisition and data analysis for mycobactin determination. ABE and BCC performed the sample acquisition, data processing and differential expression analysis for *Mtb* proteomes. KE and CB processed and sequenced RNAseq samples. AT performed the data analysis for RNAseq and all other aspects of computational analysis.

## Declaration of Interests

The authors declare no competing interests.

## References

Akashi, H., and Gojobori, T. (2002). Metabolic efficiency and amino acid composition in the proteomes of Escherichia coli and Bacillus subtilis. Proc Natl Acad Sci U S A 99, 3695–3700.

Anders, S., Pyl, P.T., and Huber, W. (2015). HTSeq--a Python framework to work with high-throughput sequencing data. Bioinformatics 31, 166–169.

Andersson, D., and Hughes, D. (2008). Effects of antibiotic resistance on bacterial fitness, viruelnce and transmission. In Evolutionary biology of bacterial and fungal pathogens, F. Baquero, C. Nombela, G.H. Cassell, and J.A. Gutierrez-Fuentes, eds. (Washington, DC: ASM Press), pp. 307–318.

Andries, K., Verhasselt, P., Guillemont, J., Gohlmann, H.W., Neefs, J.M., Winkler, H., Van Gestel, J., Timmerman, P., Zhu, M., Lee, E., et al. (2005). A diarylquinoline drug active on the ATP synthase of Mycobacterium tuberculosis. Science 307, 223–227.

Banaei-Esfahani, A., Nicod, C., Aebersold, R., and Collins, B.C. (2017). Systems proteomics approaches to study bacterial pathogens: application to Mycobacterium tuberculosis. Curr Opin Microbiol 39, 64–72.

Basan, M., Hui, S., Okano, H., Zhang, Z., Shen, Y., Williamson, J.R., and Hwa, T. (2015). Overflow metabolism in Escherichia coli results from efficient proteome allocation. Nature 528, 99– 104.

Beste, D.J., Laing, E., Bonde, B., Avignone-Rossa, C., Bushell, M.E., and McFadden, J.J. (2007). Transcriptomic analysis identifies growth rate modulation as a component of the adaptation of mycobacteria to survival inside the macrophage. J Bacteriol 189, 3969–3976.

Beste, D.J., Peters, J., Hooper, T., Avignone-Rossa, C., Bushell, M.E., and McFadden, J. (2005). Compiling a molecular inventory for Mycobacterium bovis BCG at two growth rates: evidence for growth rate-mediated regulation of ribosome biosynthesis and lipid metabolism. J Bacteriol 187, 1677–1684.

Bisson, G.P., Mehaffy, C., Broeckling, C., Prenni, J., Rifat, D., Lun, D.S., Burgos, M., Weissman, D., Karakousis, P.C., and Dobos, K. (2012). Upregulation of the phthiocerol dimycocerosate biosynthetic pathway by rifampin-resistant, rpoB mutant Mycobacterium tuberculosis. J Bacteriol 194, 6441–6452.

Borrell, S., and Gagneux, S. (2009). Infectiousness, reproductive fitness and evolution of drug-resistant Mycobacterium tuberculosis. Int J Tuberc Lung Dis 13, 1456–1466.

Borrell, S., Teo, Y., Giardina, F., Streicher, E., Klopper, M., Feldmann, J., Mueller, B., Victor, T., and Gagneux, S. (2013). Epistasis between antibiotic resistance mutations drives the evolution of extensively drug-resistant tuberculosis. Evolution, Medicine, and Public Health eot003, 65–74.

Borrell, S., and Trauner, A. (2017). Strain Diversity and the Evolution of Antibiotic Resistance. Adv Exp Med Biol 1019, 263–279.

Borrell, S., Trauner, A., Brites, D., Rigouts, L., Loiseau, C., Coscolla, M., Niemann, S., De Jong, B., Yeboah-Manu, D., Kato-Maeda, M., et al. (2018). Reference Set of Mycobacterium tuberculosis Clinical Strains: A tool for research and product development. In biorXiv (Cold Spring Harbor).

Campbell, E.A., Korzheva, N., Mustaev, A., Murakami, K., Nair, S., Goldfarb, A., and Darst, S.A. (2001). Structural mechanism for rifampicin inhibition of bacterial RNA polymerase. Cell 104, 901–912.

Casali, N., Nikolayevskyy, V., Balabanova, Y., Harris, S.R., Ignatyeva, O., Kontsevaya, I., Corander, J., Bryant, J., Parkhill, J., Nejentsev, S., et al. (2014). Evolution and transmission of drug-resistant tuberculosis in a Russian population. Nat Genet 46, 279–286.

Choi, M., Chang, C.Y., Clough, T., Broudy, D., Killeen, T., MacLean, B., and Vitek, O. (2014). MSstats: an R package for statistical analysis of quantitative mass spectrometry-based proteomic experiments. Bioinformatics 30, 2524–2526.

Comas, I., Borrell, S., Roetzer, A., Rose, G., Malla, B., Kato-Maeda, M., Galagan, J., Niemann, S., and Gagneux, S. (2012). Whole-genome sequencing of rifampicin-resistant Mycobacterium tuberculosis strains identifies compensatory mutations in RNA polymerase genes. Nat Genet 44, 106–110.

Comas, I., Chakravartti, J., Small, P.M., Galagan, J., Niemann, S., Kremer, K., Ernst, J.D., and Gagneux, S. (2010). Human T cell epitopes of Mycobacterium tuberculosis are evolutionarily hyperconserved. Nat Genet 42, 498–503.

Coscolla, M., and Gagneux, S. (2014). Consequences of genomic diversity in Mycobacterium tuberculosis. Semin Immunol 26, 431–444.

de Vos, M., Muller, B., Borrell, S., Black, P.A., van Helden, P.D., Warren, R.M., Gagneux, S., and Victor, T.C. (2013). Putative compensatory mutations in the rpoC gene of rifampin-resistant Mycobacterium tuberculosis are associated with ongoing transmission. Antimicrob Agents Chemother 57, 827–832.

du Preez, I., and Loots du, T. (2012). Altered fatty acid metabolism due to rifampicin-resistance conferring mutations in the rpoB Gene of Mycobacterium tuberculosis: mapping the potential of pharmaco-metabolomics for global health and personalized medicine. OMICS 16, 596–603.

Ehrenberg, M., Bremer, H., and Dennis, P.P. (2013). Medium-dependent control of the bacterial growth rate. Biochimie 95, 643–658.

Farhat, M.R., Shapiro, B.J., Kieser, K.J., Sultana, R., Jacobson, K.R., Victor, T.C., Warren, R.M., Streicher, E.M., Calver, A., Sloutsky, A., et al. (2013). Genomic analysis identifies targets of convergent positive selection in drug-resistant Mycobacterium tuberculosis. Nat Genet 45, 1183–1189.

Ford, C.B., Shah, R.R., Maeda, M.K., Gagneux, S., Murray, M.B., Cohen, T., Johnston, J.C., Gardy, J., Lipsitch, M., and Fortune, S.M. (2013). Mycobacterium tuberculosis mutation rate estimates from different lineages predict substantial differences in the emergence of drug-resistant tuberculosis. Nat Genet 45, 784–790.

Franzblau, S.G., Witzig, R.S., McLaughlin, J.C., Torres, P., Madico, G., Hernandez, A., Degnan, M.T., Cook, M.B., Quenzer, V.K., Ferguson, R.M., et al. (1998). Rapid, low-technology MIC determination with clinical Mycobacterium tuberculosis isolates by using the microplate Alamar Blue assay. J Clin Microbiol 36, 362–366.

Gagneux, S. (2018). Ecology and evolution of Mycobacterium tuberculosis. Nat Rev Microbiol 16, 202–213.

Gagneux, S., Long, C.D., Small, P.M., Van, T., Schoolnik, G.K., and Bohannan, B.J. (2006). The competitive cost of antibiotic resistance in Mycobacterium tuberculosis. Science 312, 1944–1946.

Gourse, R.L., Gaal, T., Bartlett, M.S., Appleman, J.A., and Ross, W. (1996). rRNA transcription and growth rate-dependent regulation of ribosome synthesis in Escherichia coli. Annu Rev Microbiol 50, 645–677.

Grandjean, L., Gilman, R.H., Martin, L., Soto, E., Castro, B., Lopez, S., Coronel, J., Castillo, E., Alarcon, V., Lopez, V., et al. (2015). Transmission of Multidrug-Resistant and Drug-Susceptible Tuberculosis within Households: A Prospective Cohort Study. PLoS Med 12, e1001843; discussion e1001843.

Gygli, S.M., Borrell, S., Trauner, A., and Gagneux, S. (2017). Antimicrobial resistance in Mycobacterium tuberculosis: mechanistic and evolutionary perspectives. FEMS Microbiol Rev.

Holmes, A.H., Moore, L.S., Sundsfjord, A., Steinbakk, M., Regmi, S., Karkey, A., Guerin, P.J., and Piddock, L.J. (2016). Understanding the mechanisms and drivers of antimicrobial resistance. Lancet 387, 176–187.

Jones, C.M., Wells, R.M., Madduri, A.V., Renfrow, M.B., Ratledge, C., Moody, D.B., and Niederweis, M. (2014). Self-poisoning of Mycobacterium tuberculosis by interrupting siderophore recycling. Proc Natl Acad Sci U S A 111, 1945–1950.

Karpinets, T.V., Greenwood, D.J., Sams, C.E., and Ammons, J.T. (2006). RNA:protein ratio of the unicellular organism as a characteristic of phosphorous and nitrogen stoichiometry and of the cellular requirement of ribosomes for protein synthesis. BMC Biol 4, 30.

Kuhl, C., Tautenhahn, R., Bottcher, C., Larson, T.R., and Neumann, S. (2012). CAMERA: an integrated strategy for compound spectra extraction and annotation of liquid chromatography/mass spectrometry data sets. Anal Chem 84, 283–289.

Lahiri, N., Shah, R.R., Layre, E., Young, D., Ford, C., Murray, M.B., Fortune, S.M., and Moody, D.B. (2016). Rifampin Resistance Mutations Are Associated with Broad Chemical Remodeling of Mycobacterium tuberculosis. J Biol Chem 291, 14248–14256.

Laxminarayan, R., Matsoso, P., Pant, S., Brower, C., Rottingen, J.A., Klugman, K., and Davies, S. (2016). Access to effective antimicrobials: a worldwide challenge. Lancet 387, 168–175.

Li, H., and Durbin, R. (2010). Fast and accurate long-read alignment with Burrows-Wheeler transform. Bioinformatics 26, 589–595.

Li, H., Handsaker, B., Wysoker, A., Fennell, T., Ruan, J., Homer, N., Marth, G., Abecasis, G., and Durbin, R. (2009). The Sequence Alignment/Map format and SAMtools. Bioinformatics 25, 2078–2079.

Love, M.I., Huber, W., and Anders, S. (2014). Moderated estimation of fold change and dispersion for RNA-seq data with DESeq2. Genome Biol 15, 550.

Luria, S.E., and Delbruck, M. (1943). Mutations of bacteria from virus sensitivity to virus resistance. Genetics 28, 491–511.

Melnyk, A.H., Wong, A., and Kassen, R. (2015). The fitness costs of antibiotic resistance mutations. Evol Appl 8, 273–283.

Minch, K.J., Rustad, T.R., Peterson, E.J., Winkler, J., Reiss, D.J., Ma, S., Hickey, M., Brabant, W., Morrison, B., Turkarslan, S., et al. (2015). The DNA-binding network of Mycobacterium tuberculosis. Nat Commun 6, 5829.

Molodtsov, V., Scharf, N.T., Stefan, M.A., Garcia, G.A., and Murakami, K.S. (2017). Structural basis for rifamycin resistance of bacterial RNA polymerase by the three most clinically important RpoB mutations found in Mycobacterium tuberculosis. Mol Microbiol 103, 1034–1045.

O’Neill, J. (2016). Review on Antimicrobial Resistance. Tackling Drug-Resistant Infections Globally: Final Report and Recommendations. J. O’Neill, ed. (London, Wellcome Trust and UK Government), p. 84.

Park, H.D., Guinn, K.M., Harrell, M.I., Liao, R., Voskuil, M.I., Tompa, M., Schoolnik, G.K., and Sherman, D.R. (2003). Rv3133c/dosR is a transcription factor that mediates the hypoxic response of Mycobacterium tuberculosis. Mol Microbiol 48, 833–843.

Peterson, E.J., Reiss, D.J., Turkarslan, S., Minch, K.J., Rustad, T., Plaisier, C.L., Longabaugh, W.J., Sherman, D.R., and Baliga, N.S. (2014). A high-resolution network model for global gene regulation in Mycobacterium tuberculosis. Nucleic Acids Res 42, 11291–11303.

Qi, Q., Preston, G.M., and MacLean, R.C. (2014). Linking system-wide impacts of RNA polymerase mutations to the fitness cost of rifampin resistance in Pseudomonas aeruginosa. MBio 5, e01562.

Reddy, P.V., Puri, R.V., Chauhan, P., Kar, R., Rohilla, A., Khera, A., and Tyagi, A.K. (2013). Disruption of mycobactin biosynthesis leads to attenuation of Mycobacterium tuberculosis for growth and virulence. J Infect Dis 208, 1255–1265.

Reiter, L., Claassen, M., Schrimpf, S.P., Jovanovic, M., Schmidt, A., Buhmann, J.M., Hengartner, M.O., and Aebersold, R. (2009). Protein identification false discovery rates for very large proteomics data sets generated by tandem mass spectrometry. Mol Cell Proteomics 8, 2405–2417.

Reiter, L., Rinner, O., Picotti, P., Huttenhain, R., Beck, M., Brusniak, M.Y., Hengartner, M.O., and Aebersold, R. (2011). mProphet: automated data processing and statistical validation for large-scale SRM experiments. Nat Methods 8, 430–435.

Reynolds, M.G. (2000). Compensatory evolution in rifampin-resistant Escherichia coli. Genetics 156, 1471–1481.

Rodriguez, G.M., Voskuil, M.I., Gold, B., Schoolnik, G.K., and Smith, I. (2002). ideR, An essential gene in Mycobacterium tuberculosis: role of IdeR in iron-dependent gene expression, iron metabolism, and oxidative stress response. Infect Immun 70, 3371–3381.

Rosenberger, G., Ludwig, C., Rost, H.L., Aebersold, R., and Malmstrom, L. (2014). aLFQ: an R-package for estimating absolute protein quantities from label-free LC-MS/MS proteomics data. Bioinformatics 30, 2511–2513.

Rost, H.L., Rosenberger, G., Navarro, P., Gillet, L., Miladinovic, S.M., Schubert, O.T., Wolski, W., Collins, B.C., Malmstrom, J., Malmstrom, L., et al. (2014). OpenSWATH enables automated, targeted analysis of data-independent acquisition MS data. Nat Biotechnol 32, 219–223.

Rost, H.L., Sachsenberg, T., Aiche, S., Bielow, C., Weisser, H., Aicheler, F., Andreotti, S., Ehrlich, H.C., Gutenbrunner, P., Kenar, E., et al. (2016). OpenMS: a flexible open-source software platform for mass spectrometry data analysis. Nat Methods 13, 741–748.

Rustad, T.R., Minch, K.J., Ma, S., Winkler, J.K., Hobbs, S., Hickey, M., Brabant, W., Turkarslan, S., Price, N.D., Baliga, N.S., et al. (2014). Mapping and manipulating the Mycobacterium tuberculosis transcriptome using a transcription factor overexpression-derived regulatory network. Genome Biol 15, 502.

Sarkar, S., Ma, W.T., and Sandri, G.H. (1992). On fluctuation analysis: a new, simple and efficient method for computing the expected number of mutants. Genetica 85, 173–179.

Schubert, O.T., Ludwig, C., Kogadeeva, M., Zimmermann, M., Rosenberger, G., Gengenbacher, M., Gillet, L.C., Collins, B.C., Rost, H.L., Kaufmann, S.H., et al. (2015). Absolute Proteome Composition and Dynamics during Dormancy and Resuscitation of Mycobacterium tuberculosis. Cell host & microbe 18, 96–108.

Schubert, O.T., Mouritsen, J., Ludwig, C., Rost, H.L., Rosenberger, G., Arthur, P.K., Claassen, M., Campbell, D.S., Sun, Z., Farrah, T., et al. (2013). The Mtb Proteome Library: A Resource of Assays to Quantify the Complete Proteome of Mycobacterium tuberculosis. Cell host & microbe 13, 602–612.

Scott, M., Gunderson, C.W., Mateescu, E.M., Zhang, Z., and Hwa, T. (2010). Interdependence of cell growth and gene expression: origins and consequences. Science 330, 1099–1102.

Seligmann, H. (2003). Cost-minimization of amino acid usage. J Mol Evol 56, 151–161.

Smith, C.A., Want, E.J., O’Maille, G., Abagyan, R., and Siuzdak, G. (2006). XCMS: processing mass spectrometry data for metabolite profiling using nonlinear peak alignment, matching, and identification. Anal Chem 78, 779–787.

Song, T., Park, Y., Shamputa, I.C., Seo, S., Lee, S.Y., Jeon, H.S., Choi, H., Lee, M., Glynne, R.J., Barnes, S.W., et al. (2014). Fitness costs of rifampicin resistance in Mycobacterium tuberculosis are amplified under conditions of nutrient starvation and compensated by mutation in the beta’ subunit of RNA polymerase. Mol Microbiol 91, 1106–1119.

Stefan, M.A., Ugur, F.S., and Garcia, G.A. (2018). Source of the Fitness Defect in Rifamycin-Resistant Mycobacterium tuberculosis RNA Polymerase and the Mechanism of Compensation by Mutations in the beta’ Subunit. Antimicrob Agents Chemother 62.

Thiele, I., Jamshidi, N., Fleming, R.M., and Palsson, B.O. (2009). Genome-scale reconstruction of Escherichia coli’s transcriptional and translational machinery: a knowledge base, its mathematical formulation, and its functional characterization. PLoS Comput Biol 5, e1000312.

Wells, R.M., Jones, C.M., Xi, Z., Speer, A., Danilchanka, O., Doornbos, K.S., Sun, P., Wu, F., Tian, C., and Niederweis, M. (2013). Discovery of a siderophore export system essential for virulence of Mycobacterium tuberculosis. PLoS Pathog 9, e1003120.

WHO (2017). Global Tuberculosis Report 2017. (Geneva, World Health Organization), p. 147.

Ycart, B. (2013). Fluctuation analysis: can estimates be trusted? PLoS ONE 8, e80958.

Zaneveld, J.R., McMinds, R., and Vega Thurber, R. (2017). Stress and stability: applying the Anna Karenina principle to animal microbiomes. Nat Microbiol 2, 17121.

Zhang, H., Li, D., Zhao, L., Fleming, J., Lin, N., Wang, T., Liu, Z., Li, C., Galwey, N., Deng, J., et al. (2013). Genome sequencing of 161 Mycobacterium tuberculosis isolates from China identifies genes and intergenic regions associated with drug resistance. Nat Genet 45, 1255–1260.

